# Thalamic neurons drive distinct forms of motor asymmetry that are conserved in teleost and dependent on visual evolution

**DOI:** 10.1101/2023.03.20.533538

**Authors:** Jacob Starkey, John Hageter, Rob Kozol, Kevin Emmerich, Jeff S. Mumm, Erik R. Duboué, Eric J. Horstick

**Affiliations:** West Virginia University, Department of Biology Morgantown, WV; Florida Atlantic University, Charles E. Schmidt College of Science, Boca Raton, FL; John Hopkins University, Department of Genetic Medicine, McKusick-Nathans Institute of Genetic Medicine, Baltimore, MD; John Hopkins University, The Center for Nanomedicine, Wilmer Eye Institute, Baltimore, MD; John Hopkins University, The Solomon H. Snyder Department of Neuroscience, Baltimore, MD; Johns Hopkins University, Department of Ophthalmology, Wilmer Eye Institute, Baltimore, MD; Florida Atlantic University, Harriet L. Wilkes Honors College, Jupiter, FL; West Virginia University, Department of Neuroscience Morgantown, WV

**Keywords:** asymmetry, visual behavior, evolution, non-model, teleost, cavefish, sensory dependent, zebrafish

## Abstract

Brain laterality is a prominent feature in Bilateria, where neural functions are favored in a single brain hemisphere. These hemispheric specializations are thought to improve behavioral performance and are commonly observed as sensory or motor asymmetries, such as handedness in humans. Despite its prevalence, our understanding of the neural and molecular substrates instructing functional lateralization is limited. Moreover, how functional lateralization is selected for or modulated throughout evolution is poorly understood. While comparative approaches offer a powerful tool for addressing this question, a major obstacle has been the lack of a conserved asymmetric behavior in genetically tractable organisms. Previously, we described a robust motor asymmetry in larval zebrafish. Following the loss of illumination, individuals show a persistent turning bias that is associated with search pattern behavior with underlying functional lateralization in the thalamus. This behavior permits a simple yet robust assay that can be used to address fundamental principles underlying lateralization in the brain across taxa. Here, we take a comparative approach and show that motor asymmetry is conserved across diverse larval teleost species, which have diverged over the past 200 million years. Using a combination of transgenic tools, ablation, and enucleation, we show that teleosts exhibit two distinct forms of motor asymmetry, vision-dependent and - independent. These asymmetries are directionally uncorrelated, yet dependent on the same subset of thalamic neurons. Lastly, we leverage *Astyanax* sighted and blind morphs, which show that fish with evolutionarily derived blindness lack both retinal-dependent and -independent motor asymmetries, while their sighted surface conspecifics retained both forms. Our data implicate that overlapping sensory systems and neuronal substrates drive functional lateralization in a vertebrate brain that are likely targets for selective modulation during evolution.

## Introduction

Differences between the left and right brain hemispheres are well-established and are observed for most cognitive processes^1^. A classic example is language in humans, where language-associated function is predominately localized in the left hemisphere^2, 3^. Behavioral asymmetries were long thought to be unique to humans, yet we now know that paw, hand, foot, and eye preferences are observed in nearly all bilateral species, and neural asymmetries likely underlie these behaviors^4–8^, which potentially provide a powerful behavioral to functional connection. Cross-species comparisons are a powerful approach to assess not only the conservation of behaviors but also to define how behavioral responses change according to the environment, behavioral adaptation, or genetic heterogeneity^9–12^. For many asymmetric behaviors, like handedness or eye preferences, genetics only partially instructs these behaviors^13–15^, suggesting that external forces such as organismal experience may also be a contributing factor. We hypothesize that comparative approaches might reveal insights into the external and internal interactions that drive behavioral asymmetry. Indeed, handedness is widely observed across diverse primates over millions of years of evolution, yet is manifested in highly varying levels of strength, directional preference (left or right-handed), and task association^16^. Similarly, visual asymmetries have been observed across avian and fish species, where each eye is preferred for specific tasks^8, 17, 18^. Pigeons and chickens are canonical models for visual asymmetry, choosing to use the right eye for visual discrimination tasks^19, 20^. Diverse fish species also show an asymmetric eye preference for visualizing conspecifics^21, 22^. However, many of these established paradigms are challenging to study, or use models not amendable to genetic and circuit manipulation. These examples illustrate that various forms of behavioral asymmetries exist, show a range of phenotypic expression, and are conserved in evolution; however, the selection of lateralization in evolution is incompletely understood.

Light-driven behaviors are conserved and essential for survival across species^23, 24^. Recently, we described a motor asymmetry in larval zebrafish that occurs during light-search behavior that is dependent on a subset of genetically defined thalamic neurons^25–, 27^. Following the loss of illumination, zebrafish larvae initiate a search pattern behavior that, at the population level, leads to increased turning, localized movement, and decreased displacement^25^ and is consistent with search patterns observed across species^28–31^. However, at the individual level, the left or right turn direction is persistent and represents a motor asymmetry, in which the direction is sustained through most of early larval development^26, 27^. This powerful and robust assay provides a foundational tool to reveal neural substrates underlying lateralized behaviors. Zebrafish provides a genetically tractable model to explore the basis of behavioral asymmetry in the form of a straight-forward left or right turn bias. It is unclear whether search-associated motor asymmetry is conserved across other fish species and what factors regulate and select for asymmetry in the brain. Comparative approaches have proven to be instrumental in understanding conservation of phenotypic traits; thus, we reasoned that assessing turning bias in diverse teleost taxa would be a powerful strategy for addressing these fundamental questions.

Here we show that motor asymmetry is conserved across diverse teleost species encompassing approximately 240 million years of evolution. We compare behavior across six different sighted teleost species representing four different Orders. We capitalized on the availability of *Astyanax*, which includes cavefish with evolutionarily-derived blindness as well as same-species sighted surface fish. Together, we show that the evolution of light search-dependent motor asymmetry is driven by vision. We also demonstrate a vision-independent form of motor asymmetry. Using acute blinding, we show that zebrafish and *Astyanax* sighted surface fish exhibit a retina-independent motor asymmetry that is not correlated with retina-driven motor asymmetry turn direction, establishing two distinct forms of motor asymmetry. Last, we demonstrate that retina-independent motor asymmetry is dependent on a subset of thalamic neurons, consistent with retina-driven responses^27^, suggesting a conserved neural basis. Collectively, our work establishes a robust model to study an evolutionarily conserved form of motor asymmetry and functional lateralization in the brain. Moreover, we reveal that vision is not a prerequisite for teleost motor asymmetry, but is required to select for motor asymmetry during evolution and is dependent on thalamic neurons.

## Results

### Motor asymmetry during light search is conserved in teleost evolution

To determine if light-search and motor asymmetry are conserved properties of larval teleost fish and how these behavioral features are modulated in evolution, we assessed the response to changes in illumination in five related fish species. We assessed 4-5 dpf old *Danionella cerebrum*^32–34^, 6-7 dpf old fathead minnow (*Pimephales promelas*)^35–37^, 5-6 dpf Mexican tetras (*Astyanax mexicanus*)^38, 39^, 12-14 dpf medaka (*Oryzias latipes*)^40, 41^, and 6-7 dpf Sheepshead pupfish (*Cyprinodon variegatus)*^42, 43^ to compare responses with Zebrafish (*Danio rerio, TL strain*). Across all species the timepoints used for testing corresponded with free-swimming larval stages. Danionella, fathead minnows, and zebrafish are all in the Order *Cypriniformes*, medaka belongs to *Beloniformes*, Sheepshead in *Cyprinodontiformes*, and Mexican tetras the Order *Characiformes* (Figure 1). For our analysis of Astyanax, we initially focused on sighted *Astyanax* morphs instead of both sighted and blind ones, as the presence of vision suggests they are more comparable to the other species tested. Our analysis covers approximately 240 million years of evolution, with ray-finned fish, including teleost, diverged from tetrapod lineages about 400 million years ago (Figure 1).

**Figure 1:**
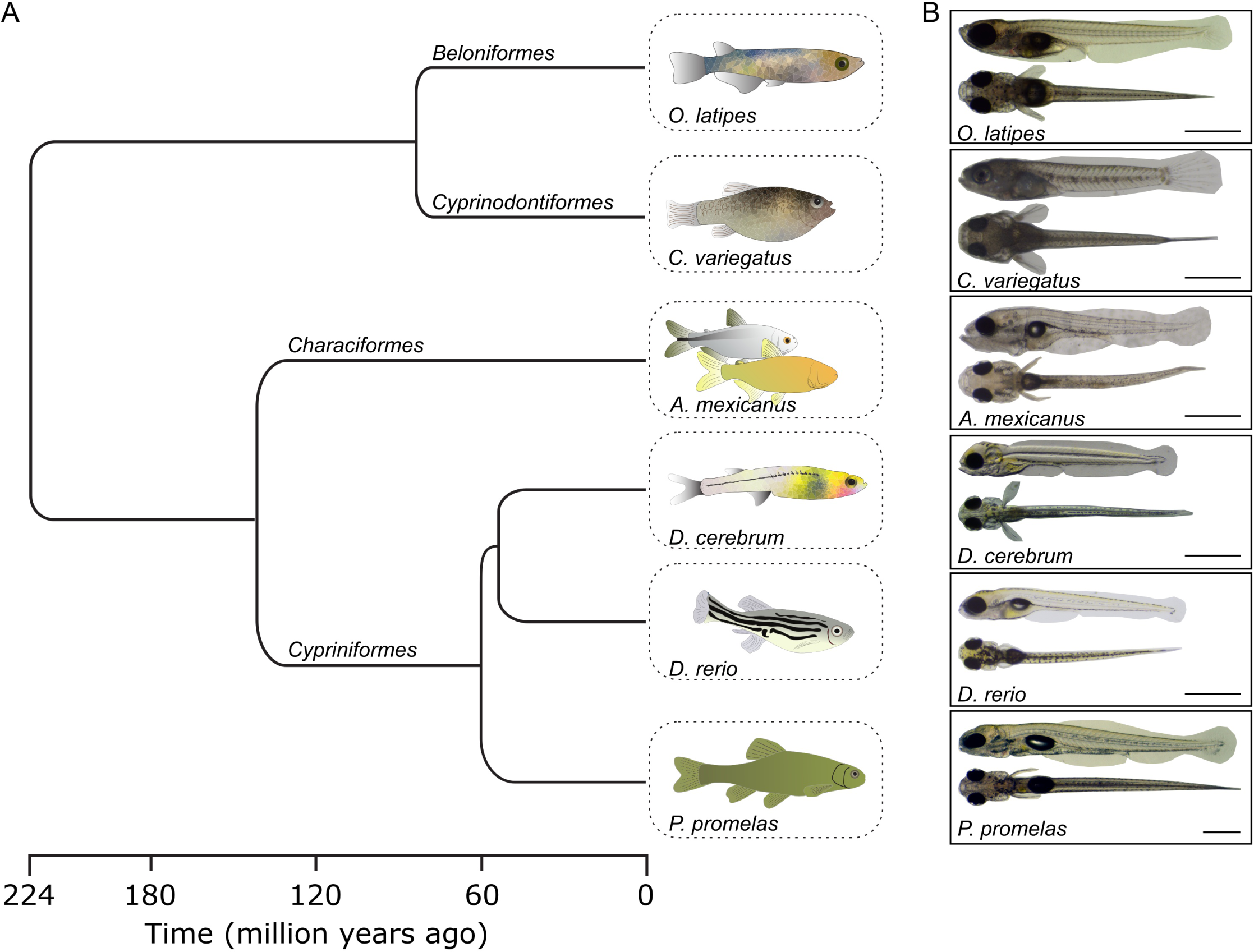
Tested teleost lineage. **(A)** Phylogenetic tree of teleost species tested. **(B)** Side and top view of the larvae for each species at the time of testing. Scale bar 1 mm.

Motor asymmetry was tested over four light transitions. Path trajectories were recorded over each of the thirty-second light transitions using custom designed behavioral apparatus (Figure 2A-B), which we have previously demonstrated provides a rigorous readout of individual left or right motor asymmetry^44^. Motor asymmetry was assessed using a ‘match index’, which is a statistical measure of the proportion of trials were turn direction matches first trial direction. At the population level, all species showed an equal distribution of left and right turn bias individuals (Figure 2C)(*D. rerio*: χ^2^(1) = 0, p=1; *P. promelas*: χ^2^(1) = 1.391, p=0.23; *D. cerebrum*: χ^2^(1) = 0.40, p=0.52; *A. mexicanus*: χ^2^(1) = 0.64, p=0.42; *C. variegatus*: χ^2^(1) = .25, p=0.62; *O. latipes*: χ^2^(1) = 0.9, p=0.34), suggesting an equal percentage of fish that turn left and right. However, within individuals, a preference for either the left or the right emerged as reflected in their higher match index following the loss of illumination (Figure 2D). We found that all teleost fish tested had strong motor asymmetry in response to the loss of environmental illumination, similar to the well-established motor asymmetry described in zebrafish (Kruskal-Wallis multiple comparisons across species: H(5) = 1.712, p=0.89)(Figure 2D). These data show that motor asymmetry is conserved across diverse sighted teleost species.

**Figure 2:**
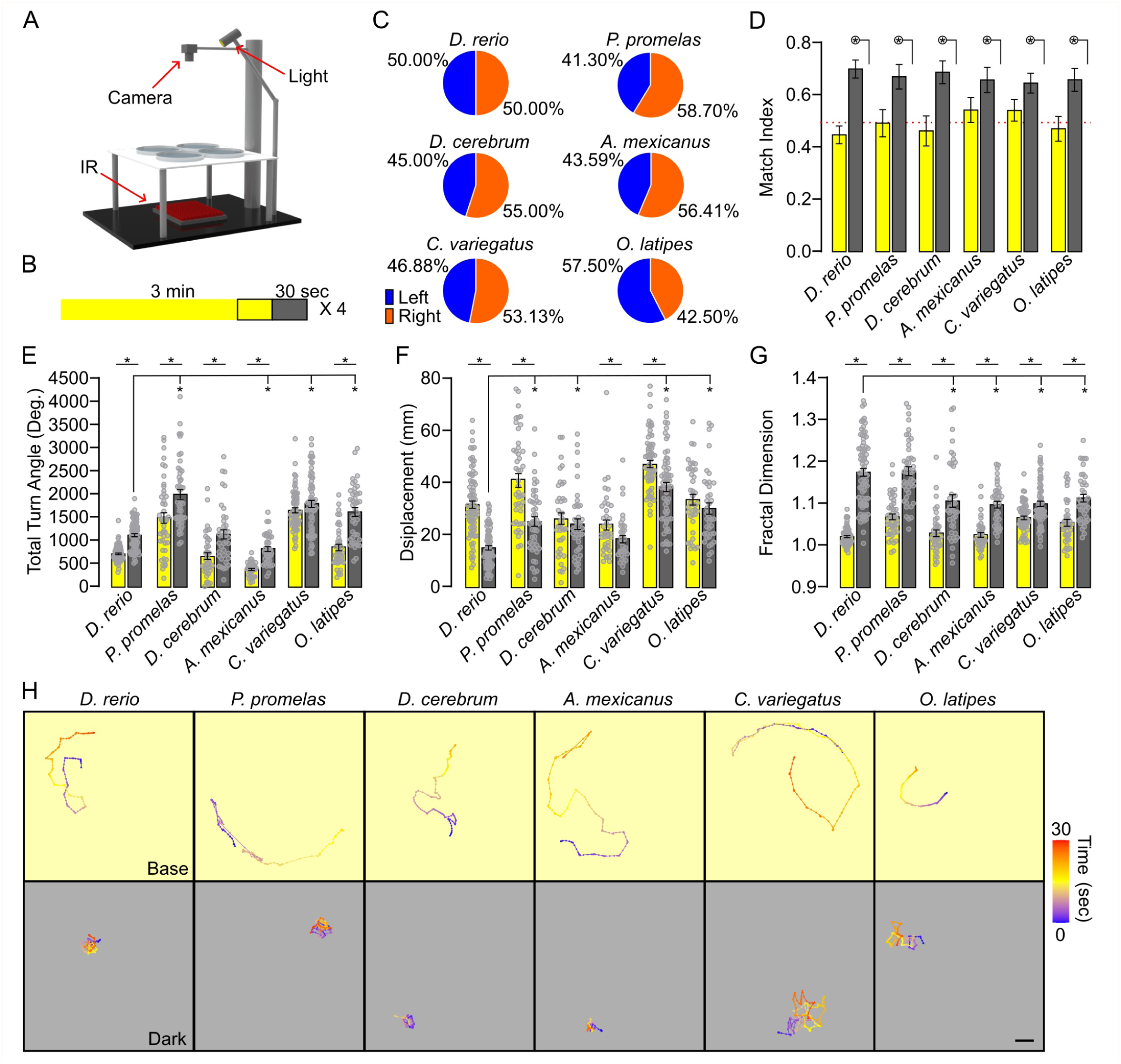
Motor asymmetry is conserved across teleost species. **(A)** Schematic for behavior testing rig and **(B)** tracking paradigm comprised showing periods of baseline illumination (yellow) and light off (gray) intervals. Outlined section indicates 30 second recording periods. **(C)** Percent of individuals showing left (blue) and right (orange) preference motor asymmetry based on the average net turn angle over all four dark recording intervals. **(D)** Match Index for *D. rerio* (N=80)*, P. promelas* (N=46)*, D. cerebrum* (N=40) *A. mexicanus* (N=39)*, C. variegatus* (N=64), and *O. latipes* (N=40) during base illumination (yellow) and dark (gray). Red dotted line at 0.5 indicates random turn bias. Circled asterisk with line indicates p<0.05 for a Wilcoxon signed rank test to 0.5. **(E-G)** Total turn angle **(E)**, displacement **(F)**, and fractal dimension **(G)** for each species. Asterisk with bar, two-tailed t-test p<0.05. **(H)** Representative trace for each species behavior during 30 second base (yellow) and dark (gray) recordings. Scale bar 10 mm. All error bars represent ±SEM.

To examine whether there were differences underlying other motor elements of search pattern behavior across species, we assessed total turning angle (TTA), displacement, and local movement (measured using a fractal dimension metric) (Figure 2E-H). In zebrafish, these parameters change in predictable manners during light search (e.g., increased turning, decreased displacement, increased local movement)^25^. The response strength of search behavior metrics varied across species (main effect based on species 1-way ANOVA *F(5,303)=33.93,* p<0.0001 (TTA); *F(5,303)=33.59,* p<0.0001 (displacement); *F(5,303)=15.91,* p<0.0001 (fractal dimension). Despite species-specific differences in search behavior, the transition from light to dark across most metrics was broadly consistent with larval zebrafish. While directional turning was observed in all fish species tested, there were subtle differences among parameters. To determine whether this due to genetic differences rather than motor asymmetry, we assessed 5 different zebrafish species. These data showed that even among zebrafish lines, subtle differences emerged, suggesting that while variations in turning are found, motor asymmetry is a generalizable feature to sighted teleosts.

### Visual input masks sensory-independent pathways driving motor asymmetry

Our previous studies in zebrafish show that photic changes are detected by the retina to instruct motor asymmetry direction^27^. However, bilaterally enucleated or genetically blind individuals, eliminating retinal drive, still exhibited motor asymmetry. These data suggest a retinal-independent pathway that also informs motor asymmetry^26^. Therefore, we next sought to address if retina-dependent and -independent motor asymmetries were related (same direction) or divergent (different directions) by testing individual larvae. We posit that a divergent form of motor asymmetry is the most parsimonious explanation to explain how sighted zebrafish could show motor asymmetry regardless of retinal input. We addressed this question using inducible transgenic tools to assay turn bias before and after the loss of visual input in individual larvae (Figure 3A). We used zebrafish lines in which retinal ganglia cells (RGCs) express a highly efficient variant of nitroreductase (NTR2.0), which allows rapid and robust ablation of cells following exposure to pro-drug metronidazole^45^. Larval fish were bred by crossing *Tg(islet2b:Gal4)*^46^ fish with *Tg(UAS:YFPcaax-v2a-NTR2.0)* breeders to drive the expression of NTR2.0 in RGCs. Before RGC ablation, groups of larvae harboring the transgene, as well as transgene-negative controls were tested for turn bias, and scored as left or right turning types (Supplementary Figure 2A – pre-ablation). Following a 24-hour period of metronidazole treatment, the RGC neuropil in the optic tectum, which is the primary visual processing center in teleost^47^, was abolished (Figure 3B). Loss of vision was confirmed by the expansion of melanophores, which occurs in the absence of visual input^48, 49^ (Figure 3C). Moreover, total turning was reduced post-ablation consistent with absent retinal input and search motor patterns^25^ (Supplementary Figure 2B). Transgene negative controls maintained turn bias direction post-ablation (Figure 3D-control). Conversely, turn bias was not maintained following RGC ablation where visual input was removed (Figure 3D-ablated). The loss of turn bias observed in genetically blinded individuals was inconsistent with our prior studies examining blind larvae^26^. Therefore, we also examined match index (MI), an agnostic indicator of motor asymmetry, which is not based on pre-ablation turn bias direction. Interestingly, we found that although blind larvae did not maintain pre-ablation turn bias direction like control larvae, motor asymmetry direction was observed, yet direction is re-established (Control: MI dark = 0.689±0.031, 1-sample Wilcoxon signed rank test to 0.5, p<0.0001; Ablated: MI dark = 0.625±0.026, 1-sample Wilcoxon signed rank test to 0.5, p<0.0001) (Figure 3E). This data suggests that loss of visual input does not abolish motor asymmetry yet unmasks a retina-independent pathway driving motor asymmetry. Taken together, motor asymmetries are likely caused by two distinct mechanisms, one that is dependent on visual input from the retina, and an extra-ocular source of photic cues.

**Figure 3:**
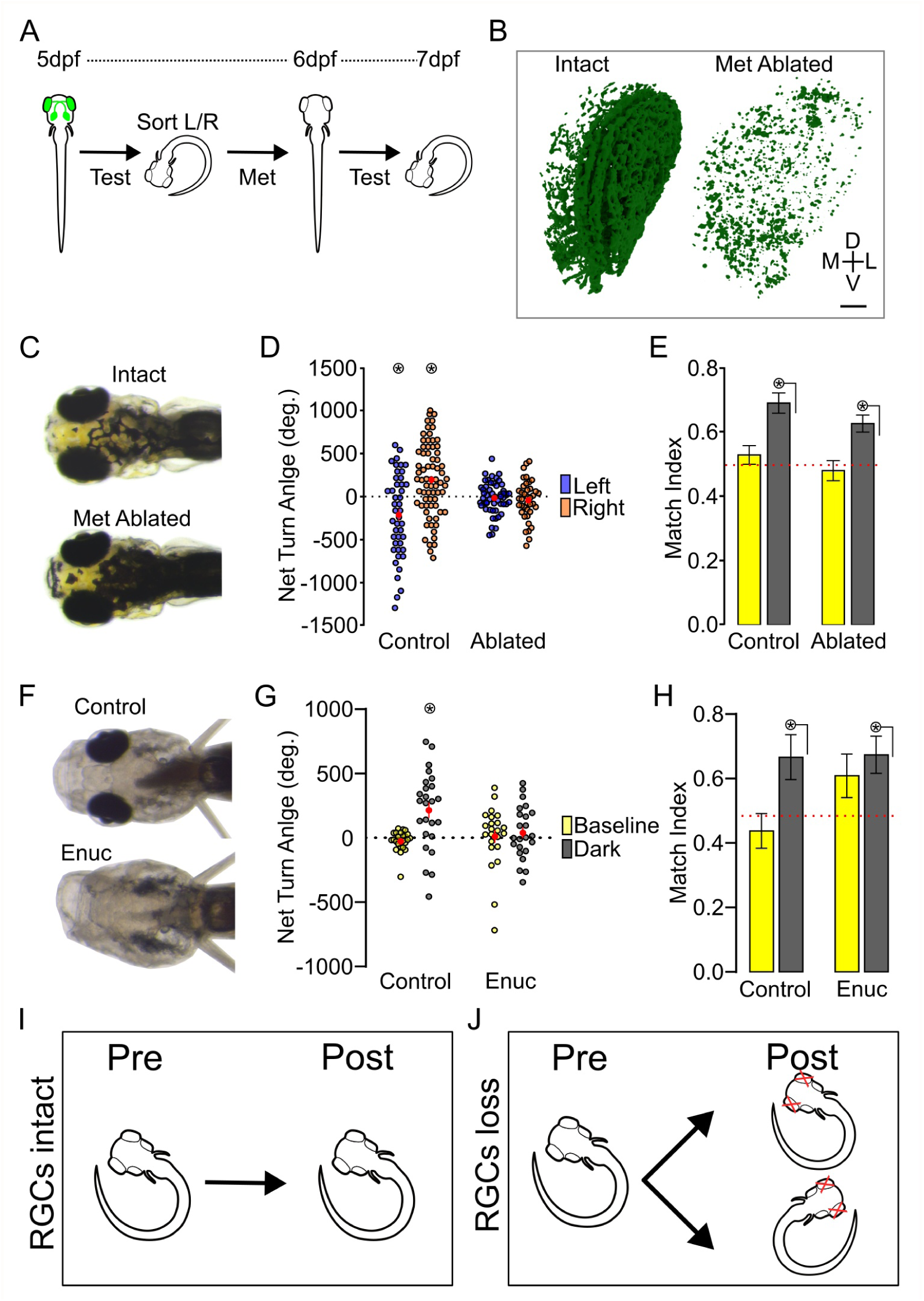
Distinct forms of motor asymmetry exist in sighted teleost. **(A)** Diagram of experimental strategy. **(B)** Illustrative of unablated intact control and ablated RGC neuropil (xxxx) in the tectum, from a single hemisphere. Scale bar 20 µm. **(C)** Melanocyte expansion in ablated blind larva compared to sighted controls. **(D)** Average individual NTA 1-day post ablation for control or ablated individuals. Circled asterisk p<0.05 one-tailed t-test to 0. (Control: Left N=45 Right N=71; Ablated: Left N=53, Right N=39). **(E)** Match index 1-day post ablation for baseline (yellow) and dark (grey) responses. Circled asterisk with line p<0.05 one-sample Wilcoxon Rank Sum test to 0.50. (Control, N=116; Ablated, N=92). Left and right groups combined **(F)** Representative *Astyanax* surface fish control (top) and enucleated (bottom). **(G)** Average NTA during baseline (yellow) and dark (gray) trials for surface fish controls (N=24) and enucleated individuals (N=23). Circled asterisk p<0.05 one-tailed t-test to 0. **(H)** Match Index for control and enucleated surface fish. Circled asterisk with line p<0.05 one-sample Wilcoxon Rank Sum test to 0.50. **(I-J)** Schematic of retina dependent and independent motor asymmetry based on retinal input **(I)** or absence **(J)**. All error bars represent ±SEM.

The discovery of an underlying motor asymmetry independent of RGC input in zebrafish leads us to inquire if this pathway is also evolutionarily conserved. To address this question, we focused on *Astyanax* surface fish, which exhibit retina-driven motor asymmetry and diverged from zebrafish approximately 120 mya (see Figure 1). Due to the lack of transgenic tools for RGC chemogenetic ablation, we used enucleation to remove visual input, which reduced total turning and confirmed absent visual drive (Figure 3F, Supplementary Figure 2C-D). Like zebrafish, we find that loss of visual input does not abolish motor asymmetry, yet it re-establishes direction (Control: MI dark = 0.667±0.070, 1-sample Wilcoxon signed rank test to 0.5, p=0.0336; Enucleation: MI dark = 0.674±0.058, 1-sample Wilcoxon signed rank test to 0.5, p=0.012)(Figure 3G-H). Therefore, both retinal dependent and independent mechanisms for maintaining persistent motor asymmetry are likely conserved in teleosts adapted to illuminated environments (Figure 3I-J).

### Motor asymmetry is loss in teleost with evolutionarily derived blindness

Search strategies are necessary to identify food, shelter, and mates^28, 29, 50^. Such behavioral strategies likely emerge in evolution as organisms develop sensory systems to assess their environments. We hypothesized that the light search behavior observed in larval fish is required to maintain animals in conditions conducive to the utilization of their senses, particularly vision for sighted species. For example, zebrafish are obligate predators by larval stages, and predatory success is dependent on vision^51^. Therefore, light search is likely employed to find and navigate to light, and we show diverse teleost larva share these motor patterns. This observation predicts that a fish that has lost its eyesight may either lose or change its responsiveness to light. Interestingly, our prior work shows that blind zebrafish maintain motor asymmetry, although light-search motor patterns are largely reduced^25, 26^. In addition, in this study we establish that retina-independent pathways also drive motor asymmetry. Therefore, it is unclear whether light as a selective pressure would or would not keep motor asymmetry. The Mexican tetra, *Astyanax mexicanus,* is a powerful model for examining naturally occurring eye loss. In addition to sighted fish, there are at least 30 populations of blind albino cavefish, which have evolved in perpetual darkness for approximately 20,000 to 1 million years^52, 53^ (Figure 4A, Supplementary Figure 3A). Despite evolutionary adaptation to a tonic dark environment, prior studies have demonstrated that cavefish maintain some forms of photo-mediated behaviors^54, 55^.

**Figure 4:**
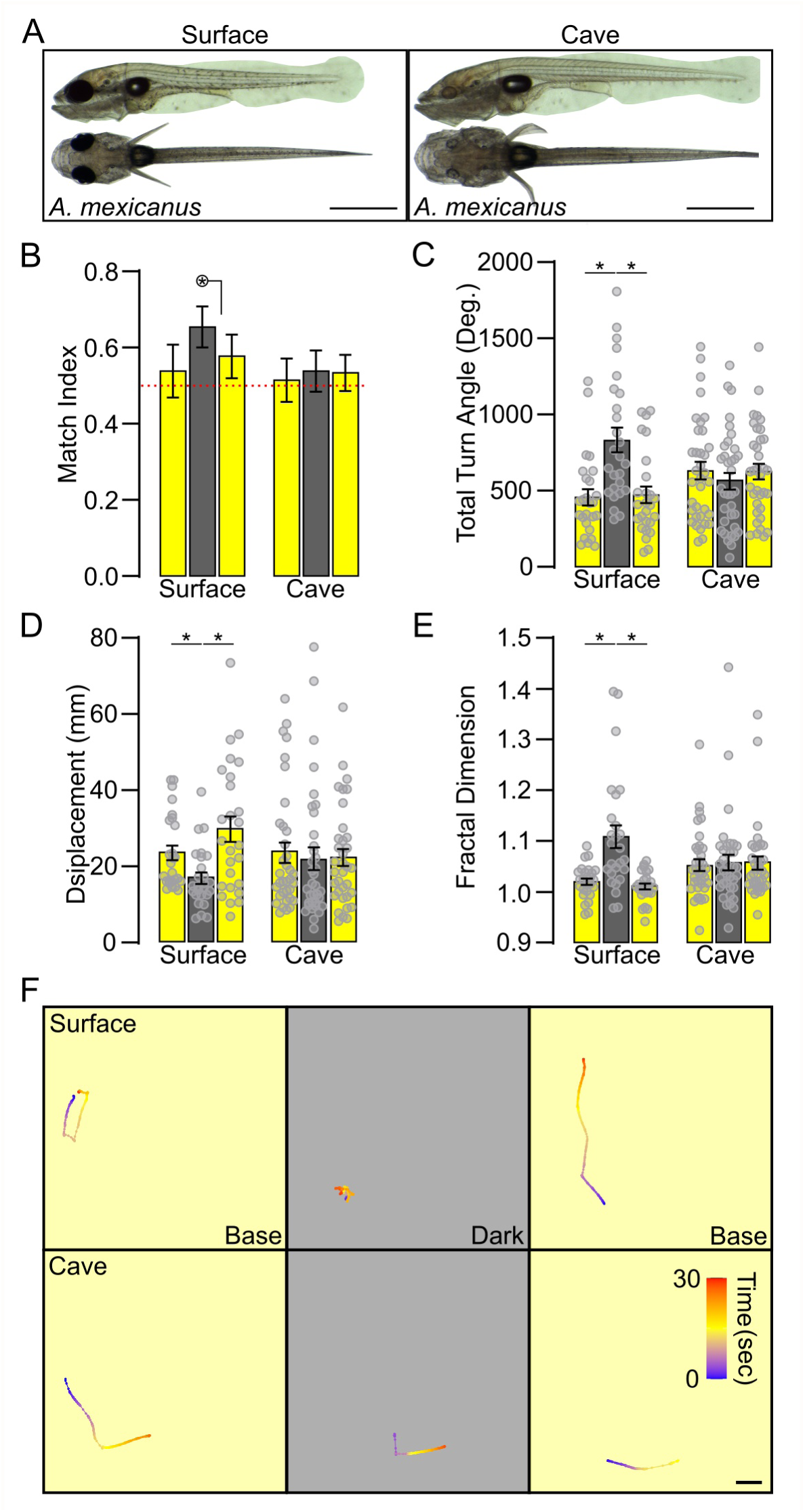
Motor asymmetry is loss in cavefish. **(A)** Representative lateral and dorsal images for surface (left) and cave (right) *A. mexicanus* at 6dpf. **(B)** Match Index for surface (N=26) and cave (N=35) populations of *A. mexicanus*. Circled asterisk with line indicates p<0.05 for a Wilcoxon signed rank test to 0.5. **(C-E)** Same as above for total turn angle **(C)**, displacement **(D)**, and fractal dimension **(E)**. Asterisk with bar, p<0.05 between groups. **(F)** Representative traces for surface (top) and cave (bottom) fish during consecutive baseline (yellow), dark (gray), and baseline following light restoration recordings. Scale bar=10mm. All error bars represent ±SEM.

We next asked whether blind cavefish had motor asymmetry comparable to other teleost species studied. Blind cavefish from the Pachón cave population and their surface conspecifics were bred by standard laboratory breeders. Each *Astyanax* surface and cavefish morph were subjected to the same assay used above with a slight modification: to control for the fact that cavefish evolved in perpetual darkness, we added another 30-second recording following the restoration of environmental illumination (Supplementary Figure 3B). As before, we assessed both individual motor asymmetry and search behavior metrics. Surface fish served as a control, and consistent with earlier results, showed robust light motor asymmetry and light-OFF responses consistent with search behavior (Figure 4B-F). Persistent same-direction turning in surface fish was also abated by light restoration, consistent with previously observed patterns in zebrafish^25^. By contrast, no observable change in match index was seen in blind cavefish, indicating an absence of motor asymmetry (light-search MI = 0.538±0.054, 1-sample Wilcoxon signed rank test to 0.5, p=0.44). Nor did we observe motor asymmetry or search patterns for a ‘dark search’ following the dark-to-light transition (dark search MI = 0.533±0.048, 1-sample Wilcoxon signed rank test to 0.5, p=0.54) (Figure 4B-F). Despite cavefish maintaining some photo-mediated behaviors and retina-independent mechanisms instructing motor asymmetry, our data shows that motor asymmetry was lost since diverging from surface fish and adapting to a perpetually dark environment.

### Thalamic input maintains sensory-independent motor asymmetry

Next, we wanted to determine the neural basis of retina-independent motor asymmetry. Previously, we demonstrated that a subset of GABAergic asymmetry-maintaining neurons (AMNs) in the thalamus maintain individual zebrafish motor asymmetry, specifically the retina-dependent mechanisms. Loss of AMNs disrupts persistent motor asymmetry, yet not overall light-search behavior, implicating a specific role in instructing motor asymmetry^26^. Moreover, AMNs show hemispheric differences in light-mediated activity that correlates with turn bias direction^27^. Our prior experiments establish that thalamic AMNs are critical to the retina-dependent motor asymmetry circuit. Therefore, we wanted to determine if AMNs were also required to maintain retina-independent motor asymmetry. To address this question, we used zebrafish and unilaterally ablated AMNs in sighted versus blind zebrafish larvae. In sighted zebrafish, two-photon ablation of AMNs, labeled by the *Tg(y279:Gal4)* driver, in a single hemisphere dictates motor asymmetry. Unilateral ablation imposes a turn bias ipsilateral to the intact hemisphere^26^, which we utilize as a control. To incorporate two-photon ablation and avoid excessive handling, we used a well-established morpholino (MO) to knockdown the transcription factor *atoh7*, which prevents RGC migration out of the retina to block all visual input to the brain^56^. Blindness in zebrafish larvae results in extensive melanophore expansion^48, 49^, which we observe in our *atoh7* MO knockdown larvae yet not control MO siblings, supporting a complete loss of photic input to the brain (Figure 5A-C). As a control, we inject scrambled standard control (SC) MO into siblings. In sighted control larvae, unilateral ablation of AMNs drives a robust contralateral motor asymmetry toward the intact hemisphere (1-sample t-test to 0, SC left ablated t(16)=9.827, p<0.0001; SC right ablated t(14)=6.942, p<0.0001) (Figure 5D), consistent with previous reports^26^. In blind larvae, unilateral AMN ablation also imposed a contralateral motor asymmetry despite the absence of retinal input (1-sample t-test to 0, *atoh7* left ablated t(8)=3.959, p=0.0042; *atoh7* right ablated t(15)=6.600, p<0.0001) (Figure 5D). Interestingly, only in sighted larvae did AMN ablation drive stronger same-direction turning, suggesting retina-dependent pathways have a stronger drive on motor asymmetry than retina-independent pathways (2-sample t-test dark control vs ablated: standard control t(242)=5.756, p<0.0001; *atoh7* t(242)=1.151, p>0.9999) (Figure 5E). No impact on turning direction strength was observed during baseline movements, suggesting that motor asymmetry was affected by AMN ablation specifically. Conversely, MI was not significantly increased, although it was elevated post ablation (Kruskal-Wallis post-hoc comparison dark control vs ablated: standard control Z=2.624, p=0.0521; *atoh7* Z=2.205, p=0.1649) (Figure 5F). Together, increased directional turning with unchanged MI suggests that AMN ablation does not increase the frequency of turn direction choice, yet the duration of same-direction turning in sighted larvae. Supporting this model, ablation had no effect on overall total turning following the loss of illumination (Supplementary Figure 4).

**Figure 5:**
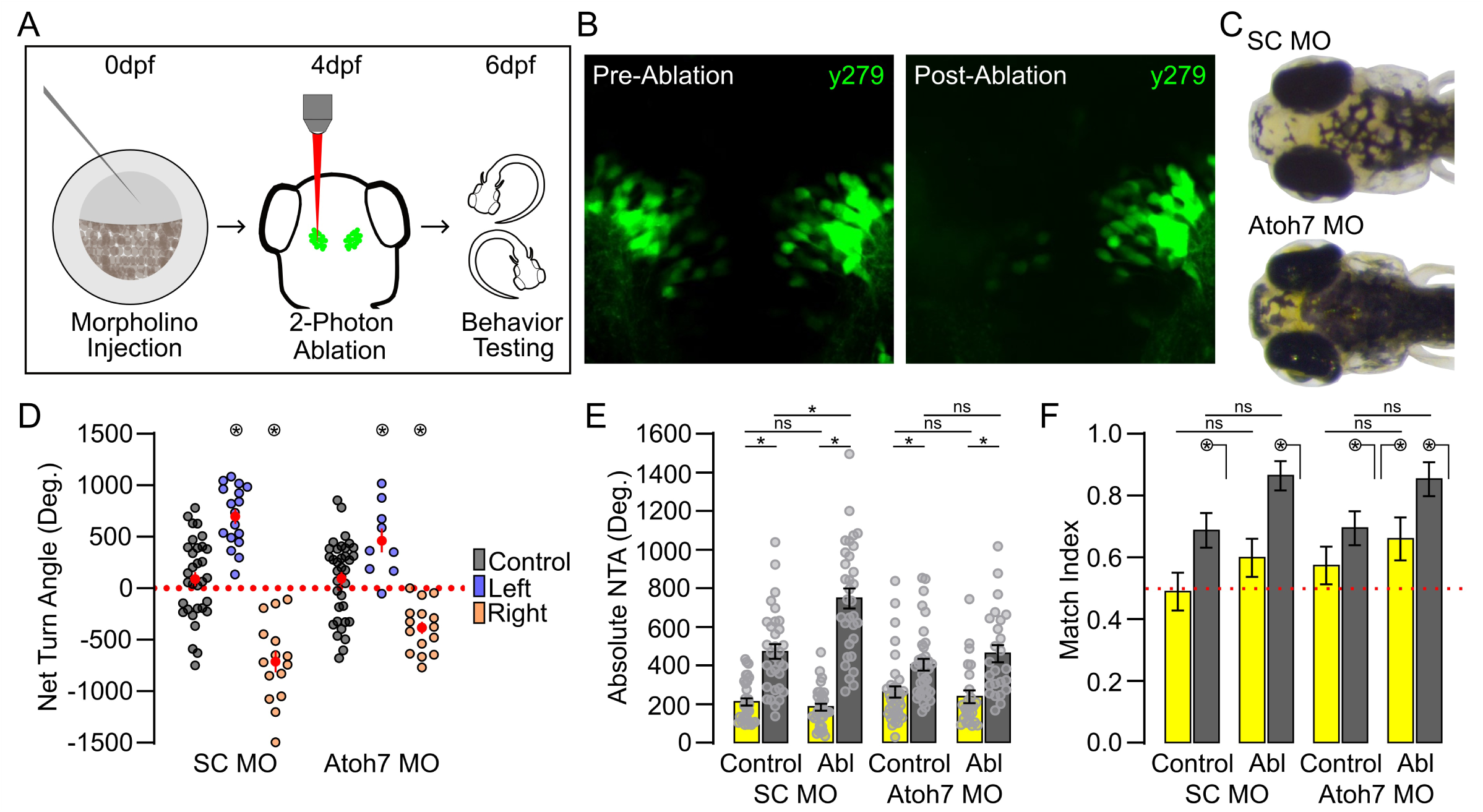
Thalamus neurons drive both forms of motor asymmetry. **(A)** Experimental timeline. **(B)** Illustrative single plane confocal scans of AMNs pre- (left) and post-ablation (right). Left hemisphere ablation shown. **(C)** Representative images of standard control (SC) MO (top) and *atoh7* MO (bottom) injected larva. **(D)** Average NTA for SC larvae (control: N=32; ablated AMNs left hemisphere N=17; ablated AMNs right hemisphere N=15) and *atoh7* MO (control: N=36; ablated AMNs left hemisphere N=9; ablated AMNs right hemisphere N=16) injected individuals. Circled asterisk p<0.05 one-tailed t-test to 0. **(E-F)** Absolute NTA **(E)** and MI **(F)** for control and ablated individuals (SC MO: Control N=32, Ablated N=32; *atoh7* MO: Control N=36, Ablated N=25). Yellow baseline illumination and grey dark path trajectories. Left and right hemisphere ablated groups combined in **E-F**. Circled asterisk with line p<0.05 one-sample Wilcoxon Rank Sum test to 0.50. Asterisk with bar, p<0.05 between groups. All error bars represent ±SEM.

In blind or sighted control larvae, which we similarly prepped yet did not perform laser ablation, no population-level motor asymmetry was observed (Figure 5D-control, grey), suggesting our experimental design was not imposing a motor asymmetry and that the observed changes were due to the loss of thalamic AMNs. Our data imply that the thalamus regulates both forms of motor asymmetry, although retina-dependent and independent mechanisms are not correlated in regards to instructing left or right turn choice. Altogether, our comparative approach suggests that the thalamus receives overlapping sensory inputs to modulate functional lateralization in the vertebrate brain and is subject to selective and potentially rapid modulation based on environmental pressures.

## Discussion

Here we take a comparative approach and show that motor asymmetry, part of a search motor pattern initially characterized in larval zebrafish, is conserved in teleost evolution. Across species with a diverse selection of traits, including cold-tolerant, slow-growing, estuarine, eury-tolerant, benthic-preferring, or pelagic-preferring, we show conservation of motor asymmetry following the loss of light. Using our comparative strategy, we could also take advantage of cavefish with evolutionarily-derived loss of vision, which exhibited no motor asymmetry. Our prior work showed that genetically blind zebrafish larvae-maintained motor asymmetry. This observation, coupled with our cavefish analysis, led us to identify that sighted-teleosts exhibit two distinct forms of motor asymmetry, retina-dependent and retina-independent. We show both forms are regulated by thalamic neurons. Here we establish a neural substrate necessary for maintaining hierarchical regulation of behavioral asymmetry and a basis for the evolution of functional and behavioral lateralization.

### Evolutionary loss of motor asymmetry in cavefish

Lateralization in the brain is proposed to improve function and behavioral performance in humans as well as other animal species^5–7, 57–59^, indicating a property that would ideally be conserved in evolution. We show that a specific form of functional lateralization, motor asymmetry, is broadly conserved in teleost. However, at first glance, the loss of motor asymmetry and related light-search motor patterns may be expected for a species adapted to a continuously dark environment. Indeed, vision and associated retinal structures have been lost in cavefish^60–63^. Moreover, crucial visual tasks in sighted teleosts, such as prey capture, were replaced in cavefish with a vibration attraction behavior (VAB) suited to navigate and forage in a cave ecosystem^64, 65^.

The loss of vision in cavefish eliminated energy-costly structures and processing, which improved fitness in a challenging environment with limited nutrient sources^66^. Nonetheless, cavefish have retained some levels of behavioral photo-responsiveness, including diurnal changes, shadow-responses, and activity within the optic tectum^54, 55, 67^. In addition, we show a retina-independent form of motor asymmetry, suggesting that motor asymmetry could be preserved in cavefish. Therefore, in the approximately 1 million years separating the surface from cave *Astyanax* populations, an evolutionarily short time period, retina-dependent and independent forms of motor asymmetry were lost. Our results indicate rapid remodeling of both retina-driven and retina-independent motor asymmetry pathways. As this remodeling occurs in such a short evolutionary time period potentially, our data suggests there was active selection against thalamic functional lateralization. Interestingly, extensive modeling using historical records suggest at least one million years is typically required for sustained divergent changes to emerge in evolution^68^. However, *Astyanax* potentially exhibits enhanced phenotypic plasticity, which could allow flexibility in behavior selection and give ancestral surface fish an advantage during their rapid colonization of cave habitats^69, 70^, and potentially account for the complete loss of motor asymmetry in such an short time scale.

### Mechanisms of retina-independent motor asymmetry

We identified that sighted teleost larvae display two distinct forms of motor asymmetry, which in a single individual are not correlated (e.g., performed in different directions). Here, we tested retina-dependent and independent turn bias over four repeated recording intervals to determine left versus right turn preference – the basis of motor asymmetry. In our prior work, we established that four trials are sufficient to rigorously characterize motor asymmetry^44^, which suggests retina-independent motor asymmetry is also a stable motor pattern and not a product of chance. Previously, we identified that a subset of asymmetry-maintaining neurons (AMNs) in the thalamus drive retina-dependent motor asymmetry and respond to visual stimuli. Moreover, visual experience during an early developmental critical period instructs functional asymmetries in the thalamus and dictates turn bias direction (retina-dependent pathway)^27^. Here, we demonstrated that this same thalamic neuron population also instructs the retina-independent pathway and turn bias direction. In our prior work, we tested thalamic responses following unilateral enucleation, which based on the structure of the teleost visual system, eliminates photic input to the contralateral hemisphere. Surprisingly, these experiments demonstrated that despite the loss of direct retinal-ganglia cell (RGC) input from the retina, not all contralateral visual responsiveness was lost in the thalamus^27^. However, unilateral enucleation would not block potential inter-hemispheric photic signaling. Regardless, retina-independent photo-responsiveness in the thalamus could be a basis of sensory-independent motor asymmetry.

What remains lacking is identifying what generates retina-independent thalamic responses to light. Extra-ocular photoreceptors (or deep brain photoreceptors) can drive photo responses in zebrafish independent of retina input^25, 71^. In zebrafish, over forty deep brain photoreceptors have been identified^72^. Therefore, one plausible hypothesis is that extra-ocular photoreception drives thalamic responses and retina-independent motor asymmetry. Alternatively, the pineal is photo-responsive and contains photoreceptors^73, 74^, which could impose retina-independent motor asymmetry. Indeed, pineal projection neurons extend to various brain regions, including the thalamus in zebrafish^75^, which is also observed in other teleosts^76^, providing a putative conserved retina-independent source of photic input to the thalamus. Unlike retina-dependent motor asymmetry, directed by asymmetric visual experience to the retina^27^, the retina-independent responses may represent a form of fluctuating asymmetry, where either pineal outputs or deep-brain photoreceptor expression in the thalamus cannot form symmetrically and impose a distinct bias in the absence of overriding retina-input.

Interestingly, cavefish retain a functional pineal and deep brain photoreceptors. Early cavefish larvae display a pineal-dependent shadow response, demonstrating that the pineal transmits signals that modulate sensorimotor responses^55^. One hypothesis to explain the lack of motor asymmetry in cavefish is a general loss of lateralization. In vertebrates, conserved molecular signaling pathways are responsible for asymmetric left-right body patterning, and in teleost near-complete leftward looping of the heart is observed^77^. Conversely, cavefish show weaker left-right patterning and a higher prevalence of reversed or symmetric heart positioning, consistent with relaxed lateralization^78^. However, our prior work demonstrated no correlation between turn bias during motor asymmetry and markers of left-right body patterning^26^. Despite weaker morphological asymmetries in cavefish, recent work shows they exhibit functionally relevant asymmetries related to VAB behavior^79^. Therefore, lateralization is still maintained in cavefish, yet the reduced morphological asymmetries and gained functional VAB asymmetries suggest these processes are subjected to active selection since diverging from sighted conspecifics. Regardless of which potential mechanism(s) drive retina-independent motor asymmetry, the absence of any apparent motor asymmetry in cavefish suggests active evolutionary selection to oppose thalamic signaling or molecular changes consistent with functional and behavioral lateralization. Moreover, our results suggest a potentially broader role of the thalamus in directing lateralization in the brain.

### Multiple forms of motor asymmetry

The thalamus instructs distinct patterns of motor asymmetry. Changing behavioral patterns are well associated with changes in the internal state, where different responses can be selected depending on internal or external cues^80–83^. Conversely, variable patterns of behavioral asymmetry are less well described. Typically, behavioral asymmetries are seen as primarily fixed. For example, handedness in humans is present early in development and is persistent^84, 85^. In the case of handedness, individuals can learn to oppose ‘natural’ handedness, yet such reversed handedness results in higher levels of brain activity, suggesting computational consequences to opposing innate or natural handedness^86, 87^. Thus, it seems unlikely that retina-independent motor asymmetry would exist as a weaker learned motor asymmetry opposing the retina-dependent form, as this would presumably exact an energetic cost. *Drosophila* exhibit an innate handedness behavior, where individuals preferentially navigate using left or right turns, dependent on the motor regulating central complex^88^. Interestingly, *Drosophila* turn bias is modulated by environmental illumination, either being reduced or directionally switched following the loss of illumination^89^. Therefore, behavioral asymmetries can be contextually modulated, although, in the case of *Drosophila*, it is unknown if turn preference in illuminated and dark conditions are both central complex dependent.

In the case of zebrafish, retina-dependent and independent are regulated by the thalamus. Retina-dependent motor asymmetry is part of a light search, and in prior studies, we have extensively discussed the likely importance of persistent same-direction turning during search behavior^44^. What remains unknown is the etiological or mechanistic importance of retina-independent motor asymmetry, which is not directionally correlated to retina-dependent behavior, suggesting it is not a secondary metric to support or enhance search motor patterns. We show that retina-independent motor asymmetry is typically masked by retina-driven responses, which likely minimizes an active etiological role. Because retina-independent motor asymmetry is only unmasked by the total removal of retina-input, suggests this change in motor asymmetry is different than the context-dependent change in turn bias asymmetry observed in *Drosophila*. We have shown that visual experience during an early critical period modulates activity patterns in the thalamus^27^. In mammals the thalamus is also a target for sensory-dependent plasticity^90–, 92^. Therefore, we hypothesize that thalamic afferent inputs, whether from the retina, other brain regions, or intrinsic deep-brain photoreceptor sensitivity, could generate functional and behavioral asymmetries due to inherent plasticity. However, these non-retinal inputs modulating thalamus function would tentatively only be observed directing motor asymmetry when stronger inputs were removed. Collectively, our work identifies that motor asymmetry is conserved in teleost and is a powerful model to characterize mechanisms that produce plasticity and lateralization in the brain.

## Methods

### Animal Husbandry

All experiments were approved by the West Virginia University Animal Care and Use Committee. All comparative zebrafish (*Danio rerio*) experiments were conducted utilizing the Tübingen long-fin (TL) strain. All zebrafish strains: Tüpfel Longfin (TL), Tübingen (Tu), AB, EkkWill (EKW), Singapore (Sing), and Wild Indian Kolkata (WIK), were raised at 28°C on a 14/10 hour light-dark cycle with a 75 µW/cm^2^ intensity light. Individuals were kept in standard E3 media ( 5mM NaCl, 0.17mM KCl, 0.33mM CaCl_2_, 0.33mM MgSO_4_-7H_2_O, and 1mM HEPES buffer pH 7.5) at a density of 40 embryos per 30 mL of E3. All experiments were conducted within the first 7 days post fertilization (dpf). Species: *Pimephales promelas, Cyprinodon variegatus, Oryzias latipes, Danionella cerebrum* (Douglass lab, University of Utah), and *Astyanax mexicanus* (Duboue lab, Florida Atlantic University) were received at 1-2 dpf. P. promelas, C. variegatus, and O. latipes were all purchased from Aquatic Research Organisms (Hampton, NH). Larvae were assessed upon arrival, and any dead, deformed, or underdeveloped individuals were removed. *D. cerebrum,* fathead minnow (*P. promelas*), and Mexican tetras (*A. mexicanus*) were raised using the standard procedure stated above until 5, 7, and 6 dpf respectively. Sheepshead pupfish (*C. variegatus*) were raised in a 5 part per thousand (ppt) salt solution at 27°C with standard light-dark cycle until 7dpf. Medaka (*O. latipes*) were raised at 25°C under standard conditions until 14-16 dpf.

### Behavior tracking and analysis

Tracking was conducted at the following ages: 4-5 dpf (*D. cerebrum)*, 5-6 dpf (*A. mexicanus*), 6-7 dpf (*Danio rerio, P. promelas, C. variegatus*), and 12-14 dpf (*Oryzias latipes*) unless otherwise stated. Larvae were tracked in a multiplexed arena made up of four 9 cm dishes. Larvae were tracked under infrared illumination (940nm, CMVision Supplies) using a µEye IDS1545LM-M CMOS camera (1^st^ Vision) and 19 mm lens (Thorlabs, MVL12WA) equipped with a 780 nm long-pass filter (Thorlabs, FGL780). Environmental illumination was supplied by a cool white LED (Thorlabs, MCWHL5) tuned to ∼55 µW/cm^2^ recorded at the stage level using an optometer (International Light Technologies, ILT2400). Behavior was recorded between 26 and 28°C during the day light cycle at least 2 hours following lights coming on and 2 hours before lights turning off. Larvae were tracked using a custom DAQtimer script operating through IDL to control the camera, lights, and tracking parameters. Tracking parameters consisted of four repeated 30 second tracking periods at 10 fps in baseline illumination and light off stimulus separated by 150 seconds of reacclimating to baseline illumination. Tracking parameters for comparison between cave and surface variants of *A. mexicanus* were conducted as stated above with an additional 30 second tracking period of baseline illumination following light off stimulus. We analyzed these trajectories using 6 measures: Net turn angle (NTA), match index (MI), absolute NTA, fractal dimension (FD), total turning angle (TTA), and displacement. Trials with less than 10 seconds of points collected during a tracking period were excluded. Individuals missing three datapoints during baseline or dark tracking were excluded. NTA is the summation of all trajectory changes during the tracking period with leftward and rightward movement having (-) or (+) sign, respectively. MI is the proportion of trials that match the direction of the first trial. TTA is the summation of the magnitude for all trajectory changes. Fractal dimension is a measure used to calculate the complexity of a trajectory, where 1 is a straight line and 2 is a line completely filling 2D space. Displacement shows the distance between the first and final points of a trial.

### Enucleations

Enucleations were performed with surface *A. mexicanus* fish. Larvae were received as described above, and raised until 4 dpf. At 4 dpf, individuals underwent behavior testing and sorted for leftward and rightward bias. Left and right categorized fish were immobilized with a small dose of tricaine (MS-222) after which both eyes were manually removed. Enucleated larvae recovered in a 1X Evans solution for 24 hours before being transitioned to E3 media on 5 dpf, and behavioral recordings were conducted at 6 dpf. Controls were handled and treated in the same manner yet without enucleation.

### Morpholino Injections

At the 1-cell stage, embryos were injected with 4pg of a morpholino against *Atoh7* target (5’-TTCATGGCTCTTCAAAAAAGTCTCC-3’) or a standard control target (5’-CCTCTTACCTCAGTTACAATTTATA-3’). To confirm injection, 120pg of RFP RNA was co-injected and used to sort larvae at 1 dpf. Loss of vision was verified through the presence of increased melanocyte development at 6-7dpf.

### Ablations

#### Genetic Ablations

For nitroreductase ablations, *Tg(isl2b:Gal4; 5xUAS:GAP-tagYFP-P2A-nsfB_Vv-F70A/F109Y)* were bred and sorted for reporter expression at 2-4 dpf. Individuals were split into two groups, transgene positive or negative, and behavior tested at 5 dpf as previously described. Larvae were grouped as left or right biased based on behavioral performance. Following behavioral classification larvae were treated overnight (20 hours) with 1mM Metronidazole in E3 embryo media to induce ablation. Ablations were confirmed by visualizing melanocyte expansion and loss of reporter fluorescence after groups were removed from Metronidazole. Larvae were maintained in fresh E3 media for 24 hours and behavior tested again at 7 dpf.

#### Laser Ablations

*Tg(y279:Gal4, UAS:Kaede)* larvae were injected with atoh7 or control morpholino as above and raised in 200 μM PTU in E3 until 4 dpf. At 4 dpf, individuals were mounted in 2% low-melt agarose for imaging and laser ablation. Larvae were imaged using a Scientifica two-photon Vivoscope with Spectra-Physics MaiTai laser and 16x water immersion objective, and image acquisition was performed through the use of MatLab ScanImage software. Representative single-plane images of AMNs pre and post-ablation were captured at a 7x zoom with a wavelength of 940nm and power of 15 μW. Ablations were conducted through multiple 2 second acquisitions at a 70x zoom and power of 80 μW. Successful ablation was determined by the loss of fluorescence, formation of local cavitation, and tissue distortion. Individuals post-ablation were raised for 24 hours in 1x Evans solution and transitioned to E3 media for 24 hours before behavior testing at 6 dpf.

## Statistical Analysis

All statistical analyses were performed in Prism (Graphpad) and RStudio (R Core Team). Data presented in figures represent means ± SEM. Non-normally distributed data was analyzed using the Kruskal-Wallis test, adjusted with the Dunn’s correction, for factorial comparisons, or Wilcoxon signed-rank tests for one- and two-way comparisons. Normally disturbed data was compared using either 1- or 2-way t-tests. For multiple comparisons, ANOVAs were performed and adjusted with a Bonferroni correction. Phylogenetic tree design and analysis was composed utilizing the public database for evolutionary timescale, TimeTree 5^93^.

## Acknowledgements

We thank the Bergeron lab for access to imaging resources to capture live images of the different species tested. We also thank the Douglass lab for sharing *Danionella* embryos. Last, we thank Sarah Ackerman for manuscript review and commentary.

## Funding

This work was supported by West Virginia University and Department of Biology startup funds, the Research and Scholarship Advancement award, Program to Stimulate Competitive Research funds, and NIGMS P20GM144230-02 provided to EJH. NIMH R15MH118625-01 and R15 MH132057-01, and NSF EDGE1923372 to ERD. NIH awards R01OD020376, and a program core grant to the Wilmer Eye Institute P30EY001765 to JSM and NIH/NRSA predoctoral fellowship award F31EY032790 to KE.

## Author Contributions

EJH conceived the experiments and wrote the manuscript with JS, JH, and ERD. ERD and RK provided *Astyanax* and expertise required to perform associated behavior experiments and manipulations. KE and JSM generated lines for chemogenetic ablation of RGCs. All authors approved the final manuscript.

## Declaration of Interests

The authors declare no competing interests

## Notes

### Competing Interest Statement

The authors have declared no competing interest.

