## Supplementary Data for "Thalamic neurons drive distinct forms of motor asymmetry that are conserved in teleost and dependent on visual evolution"

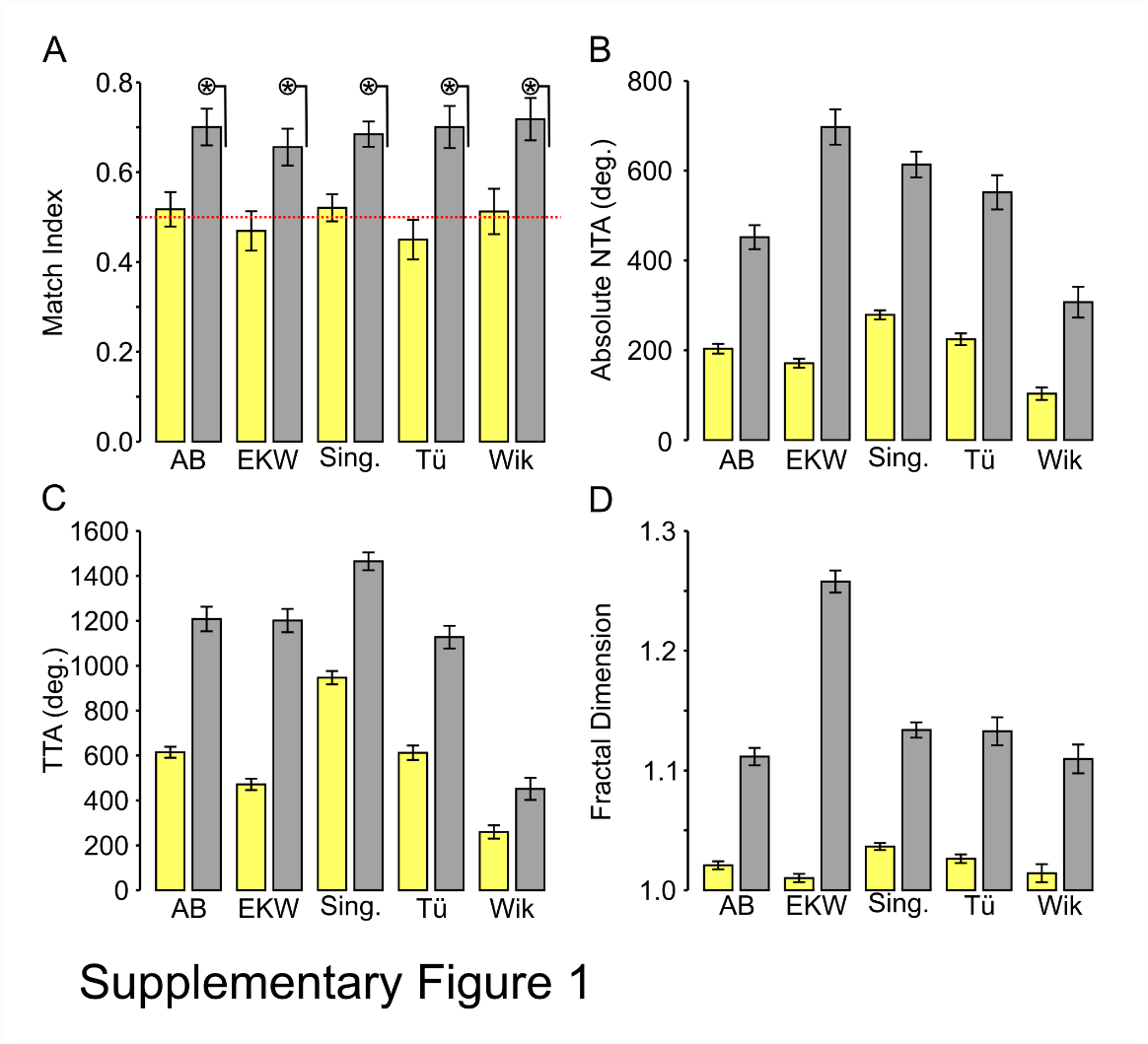

**Supplemental Figure 1: Zebrafish strains motor asymmetry and search behavior.** Measures of motor response among wildtype zebrafish strains (AB, N=50; EKW, N=59; Singapore, N=104; Tübingen, N=50; Wik, N=43) during baseline (yellow) and dark (grey). **(A)** Match index (Kruskal-Wallis Baseline: H(5)=4.28, p=0.510; Dark: H(5)=1.64, p=0.897), **(B)** Absolute average NTA (F(5,380) = 15.247, p<0.0001) **(C)** total turn angle (F*(5,380) = 34.814*, p<0.0001) and **(D)** fractal dimension (F*(5,380) = 46.331*, p<0.0001). Circled * indicates p <0.05 one sample Wilcoxon signed rank test to 0.5.

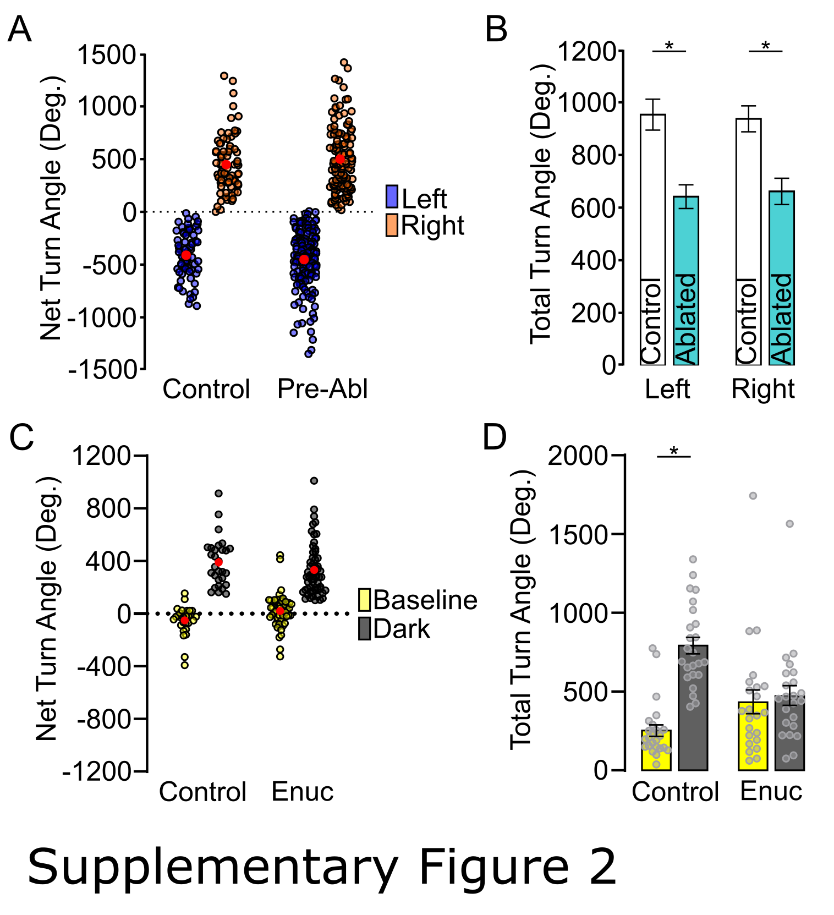

**Supplemental Figure 2: Characterization of retina dependent and independent motor asymmetry. (A)** Pre-ablation average NTA. Individuals were sorted as left biased (blue) or right biased (orange) (Control: Left N=71, Right N=78; Ablated: Left N=145, Right = 127). **(B)** Left and right turn bias larvae 1-day post ablation total turning for control (white) and ablated (blue) individuals. (Control: Left N=45, Right N=71; Ablated: Left N=53, Right = 39). **(C)** Pre-Enucleation average NTA during baseline (yellow) and dark (gray) for surface fish controls (N=29) and enucleated (N=58) larvae. Circled asterisk p<0.05 one-tailed t-test to 0. **(D)** Total turning during baseline and dark recordings for control (N=24) and enucleated (N=23) surface fish. Asterisk with bar, p<0.05 between groups. Asterisk with bar, p<0.05 between groups.

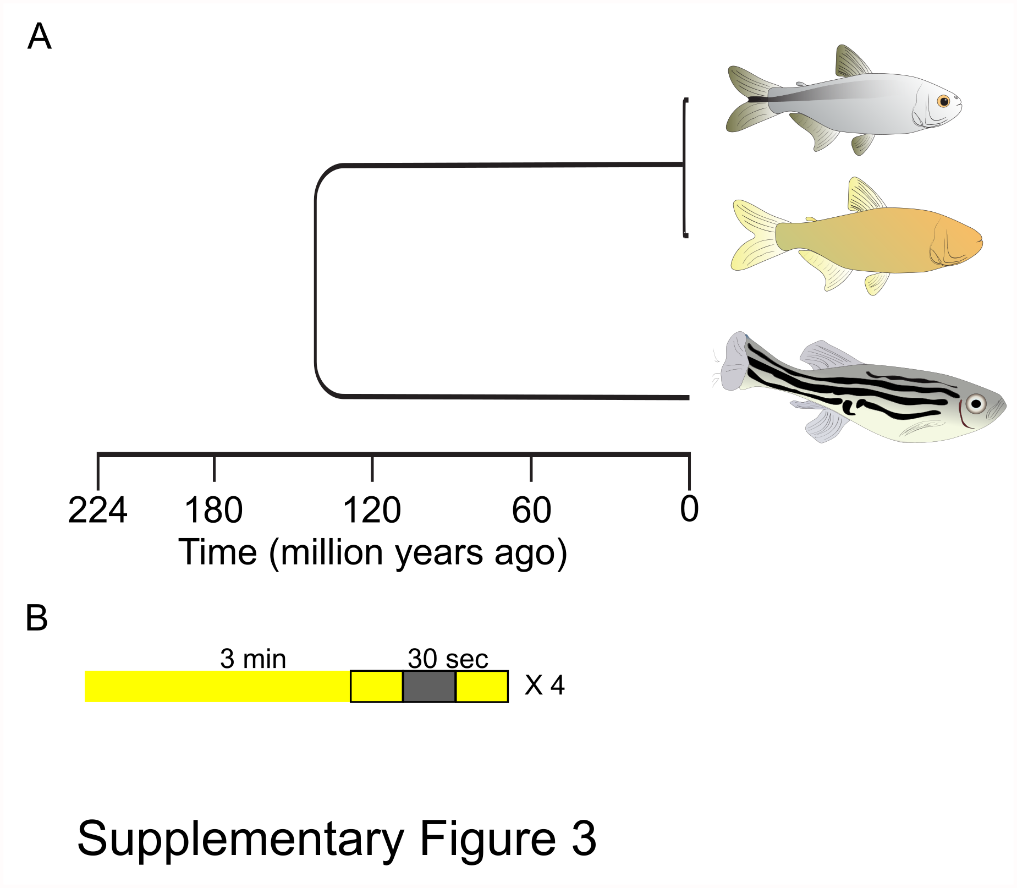

**Supplemental Figure 3: Characterization of Astyanax cavefish motor asymmetry. (A)** Phylogenetic tree for *D. rerio* (out group) and *A. mexicanus* surface and cave morphs. **(B)** Recording paradigm utilized for cave and surface fish showing periods of baseline illumination (yellow) and dark (gray) periods. Outlined sections indicate 3 consecutive 30 second recording periods.

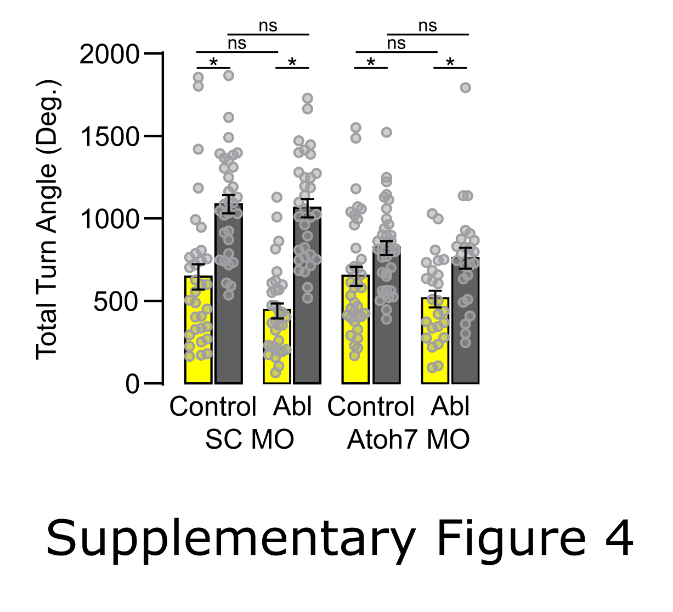

**Supplemental Figure 4: Ablations does not affect total turning.** Total turning for control and ablated individuals (SC MO: Control N=32, Ablated N=32; *atoh7* MO: Control N=36, Ablated N=25). Asterisk with bar, p<0.05 between groups.
